# Delving deeper: Relating the behaviour of a metabolic system to the properties of its components using symbolic metabolic control analysis

**DOI:** 10.1101/356139

**Authors:** Carl D. Christensen, Jan-Hendrik S. Hofmeyr, Johann M. Rohwer

## Abstract

High-level behaviour of metabolic systems results from the properties of, and interactions between, numerous molecular components. Reaching a complete understanding of metabolic behaviour based on the system’s components is therefore a difficult task. This problem can be tackled by constructing and subsequently analysing kinetic models of metabolic pathways since such models aim to capture all the relevant properties of the system components and their interactions.

Symbolic control analysis is a framework for analysing pathway models in order to reach a mechanistic understanding of their behaviour. By providing algebraic expressions for the sensitivities of system properties, such as metabolic fluxor steady-state concentrations, in terms of the properties of individual reactions it allows one to trace the high level behaviour back to these low level components. Here we apply this method to a model of pyruvate branch metabolism in *Lactococcus lactis* in order to explain a previously observed negative flux response towards an increase in substrate concentration. With this method we are able to show, first, that the sensitivity of flux towards changes in reaction rates (represented by flux control coefficients) is determined by the individual metabolic branches of the pathway, and second, how the sensitivities of individual reaction rates towards their substrates (represented by elasticity coefficients) contribute to this flux control. We also quantify the contributions of enzyme binding and mass-action to enzyme elasticity separately, which allows for an even finer-grained understanding of flux control.

These analytical tools allow us to analyse the control properties of a metabolic model and to arrive at a mechanistic understanding of the quantitative contributions of each of the enzymes to this control. Our analysis provides an example of the descriptive power of the general principles of symbolic control analysis.

**Author summary:** Metabolic networks are complex systems consisting of numerous individual molecular components. The properties of these components, together with their non-linear interactions, give rise to high-level observed behaviour of the system in which they reside. Therefore, in order to fully understand the behaviour of a metabolic system, one has to consider the properties of all of its components. The analysis of computer models that capture and represent these systems and their components simplifies this task by allowing for an easy way to isolate the effects of each individual component. In this paper we use the framework of symbolic control analysis to investigate the sensitivity of the rate of flow of matter through one of the branches in a particular metabolic pathway towards changes in the rates of individual reactions. Here we are able to quantify how certain chains of reactions, individual reactions, and even thermodynamic and kinetic aspects of individual reactions contribute to the overall sensitivity of the rate of matter-flow. Thus, we are able to trace the behaviour of the system as a whole in a mechanistic way to the properties of the individual molecular components.

## Introduction

Metabolic systems are highly complex and interconnected networks consisting of numerous functional molecular components. These systems exemplify the phenomenon of emergence, since behaviour at the system level rarely relies on any single component, arising rather from the unique properties and non-linear interactions of all its components. Unfortunately this means that understanding these systems on a mechanistic basis is a challenging task that requires quantitative knowledge of all components together with their interactions. In this regard, kinetic models of metabolic systems are essential tools [1,2], as they allow for the integration of kinetic information on the constituent enzymes and subsequent calculation of system behaviour on a scale that is otherwise difficult, if not impossible, to achieve with even the most meticulous laboratory techniques. However, metabolic models are not, in themselves, sufficient to achieve the sought-after mechanistic understanding. The size and complexity of kinetic models are approaching those of the systems they are representing (e.g. [3]). There is therefore a need for theory and tools that allow for systematic and quantitative investigation into the origins of emergent properties of the system.

The framework of metabolic control analysis (MCA) is one such tool that aims to explain the behaviour of a system in terms of its components [4,5]. This framework allows for the quantification of the sensitivity of system variables (such as fluxes and steady-state concentrations) towards perturbations in the activities of the reactions of a metabolic system. Additionally, it allows for these sensitivities to be related to the network topology and the properties of the pathway components through the application of the summation and connectivity theorems of MCA. The theory and methods that constitute MCA have been expanded upon extensively since its initial conception [6–8], and methods have been developed that generalise the summation and connectivity theorems [9]. However, MCA is frequently only applied to determine the control of certain key metabolic variables with the end goal of metabolic engineering [10,11], without considering how this control is brought about. In other words, the question of explaining emergence is often ignored in favour of more practical goals. While a more pragmatic approach is certainly understandable, it leaves the question unanswered how these control properties of metabolic systems can be understood mechanistically in terms of their components.

An extension to conventional MCA that seeks to address this shortcoming is the framework of symbolic metabolic control analysis [12,13] and the related method of control-pattern analysis [14]. In this framework symbolic (or algebraic) expressions of control coefficients, which consist of terms representing the individual steps within a metabolic pathway, are generated and analysed. Thus, in addition to ascribing a numeric value to the sensitivity of a system property towards a perturbation, this methodology provides a means for understanding how the sensitivity arises.

Another avenue that can lead to a more complete understanding of metabolic systems is through the study of their thermodynamics. In structural metabolic models for flux balance analysis, for instance, the thermodynamics of a system is already one of the criteria used to constrain the possible solution space [15]. In kinetic models, the distance of a reaction from equilibrium can indicate whether an enzyme-catalysed reaction is predominantly controlled by the properties of the enzyme itself, or by the intrinsic mass action effect [7,16–18]. However, in the past, the practical application and utility of this idea has been limited due to an imprecise delineation between what can be considered near-equilibrium and far-from-equilibrium [7, 16]. Additionally, even if distance from equilibrium is precisely defined, it does not give complete insight into the relative importance of the effect of enzyme binding as opposed to that of mass action [16].

One organism in which the systems biology approach has successfully yielded new insight is *Lactococcus lactis* (reviewed in [19]). Much work has specifically gone into understanding glycolysis in this organism, as is evident from the variety of published models that describe this system [20–23]. However, the exact mechanism behind the switch between mixed-acid fermentation and the lower ATP-yielding homolactic fermentation is yet to be uncovered [19]. Previous work in this regard has pointed to redox balance as playing an important role [24–28]. As part of a larger study that focussed on the quantification of regulatory routes in metabolic models [29], we utilised generalised supply-demand analysis [30] to uncover the effect of the redox balance on the different metabolic branches of pyruvate metabolism of *Lactococcus lactis* using a previously published metabolic model [21]. In that study, an increase in NADH/NAD^+^ was shown to decrease flux towards acetaldehyde and ethanol, mirroring past experimental [24–27] and FBA modelling [28] findings. This phenomenon was shown to originate predominantly from the interaction of NADH/NAD^+^ with pyruvate dehydrogenase. Additionally, we quantified this effect in terms of control and elasticity coefficients, thereby distinguishing between the contribution of systemic and local effects to the observed flux response.

In this paper we build on the above-mentioned work by examining the origin of the control of pyruvate dehydrogenase on the flux through the acetaldehyde dehydrogenase-catalysed reaction in *L. lactis*. To this end we employ the methods of symbolic control analysis and thermodynamic/kinetic analysis. First, we consider algebraic expressions of control coefficients in terms of elasticity coefficients and fluxes [12,13]. These symbolic expressions are examined within the framework of control-pattern analysis [14]. By identifying, quantifying, and comparing common motifs within control patterns we determine the importance of different chains of local effects over a range of NADH/NAD^+^ values, thus explaining metabolic control in physiological terms. Second, we consider how the elasticity coefficients that make up the control coefficient expressions change with NADH/NAD^+^, in particular with regard to the contribution of enzyme binding and of mass-action. By relating these results to one another we show how the properties of the individual reactions lead to the observed control profile. In this way we not only explore the properties of the system in question, but also attempt to demonstrate general principles for understanding metabolic systems within the context of these frameworks.

## Materials and methods

### Metabolic control analysis

Metabolic control analysis (MCA) is a framework for quantifying the control properties of a steady-state metabolic system in terms of the responses of its fluxes and metabolite concentrations towards perturbations in the rates of its reactions [4, 5]. Below we briefly describe the fundamental coefficients of MCA and define their relationships. For a more complete treatment of these concepts see [7].

The elasticity coefficients of a reaction describe the sensitivity of its rate towards perturbations in its parameters or the concentrations of its substrates, products, or direct effectors (i.e., its variables) in isolation. An elasticity coefficient 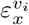 denotes the ratio of the relative change of the rate *v*_*i*_ of reaction *i* to the relative change in the value of parameter or concentration *x*. Elasticity coefficients are local properties of the reactions themselves.

A control coefficient describes the sensitivity of a system variable of a metabolic pathway (e.g., a flux or steady-state metabolite concentration) towards a perturbation in the local rate of a pathway reaction. The control coefficient 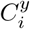 denotes the ratio of the relative change of system variable *y* to the relative change in the activity of reaction *i*. Unlike elasticity coefficients, control coefficients are functions of the complete system and depend on both network topology, as well as the properties of pathway components and their interactions.

MCA relates the above-mentioned sensitivities to the properties of the system components through summation and connectivity properties [4]. The flux-summation property, which describes the distribution of control between reactions within pathway, states that the sum of the control coefficients of all reactions on any particular flux is equal to 1. The flux-connectivity theorem describes the relationship between flux-control and elasticity coefficients. It states that when a metabolite affects multiple reactions, the sum of the products of each of the control coefficients of a particular flux with respect to these reactions multiplied by their corresponding elasticity coefficients is equal to 0.

For example, for the two-step system with reactions 1 and 2,

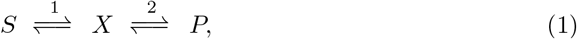

solving the summation 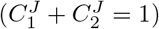 and connectivity 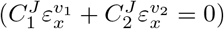 equations simultaneously produces expressions for the two control coefficients 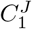 and 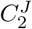 in terms of the elasticity coefficients:

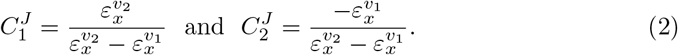

Summation and connectivity relationships also exist for concentration control coefficients, but they will not be treated here since they do not enter the present analysis.

### Symbolic metabolic control analysis

The relationship between control and elasticity coefficients expressed above can also be obtained through one of the various matrix-based formalisations of MCA [9,31–37]. One such method combines the summation and connectivity properties into a generalised matrix form called the control matrix equation 9. Here a matrix of independent control coefficients, **C**^**i**^, and a matrix expressing structural properties and elasticity coefficients, **E**, are related by

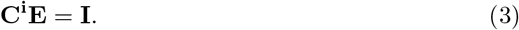

**C**^**i**^ can therefore be calculated by inverting **E**. Algebraic inversion of **E** yields expressions similar to those shown in Eq. 2 [12].

Symbolic control analysis is based on the idea that algebraic control coefficient expressions also translate into physical concepts. Each term of the numerator of a control coefficient expression represents a chain of local effects that radiates from the modulated reaction throughout the pathway as demonstrated in Fig. 1 [14]. These chains of local effects, known as *control patterns,* describe the different paths through which a perturbation can affect a system variable, thereby partitioning the control coefficient into a set of additive terms.

**Fig 1.**
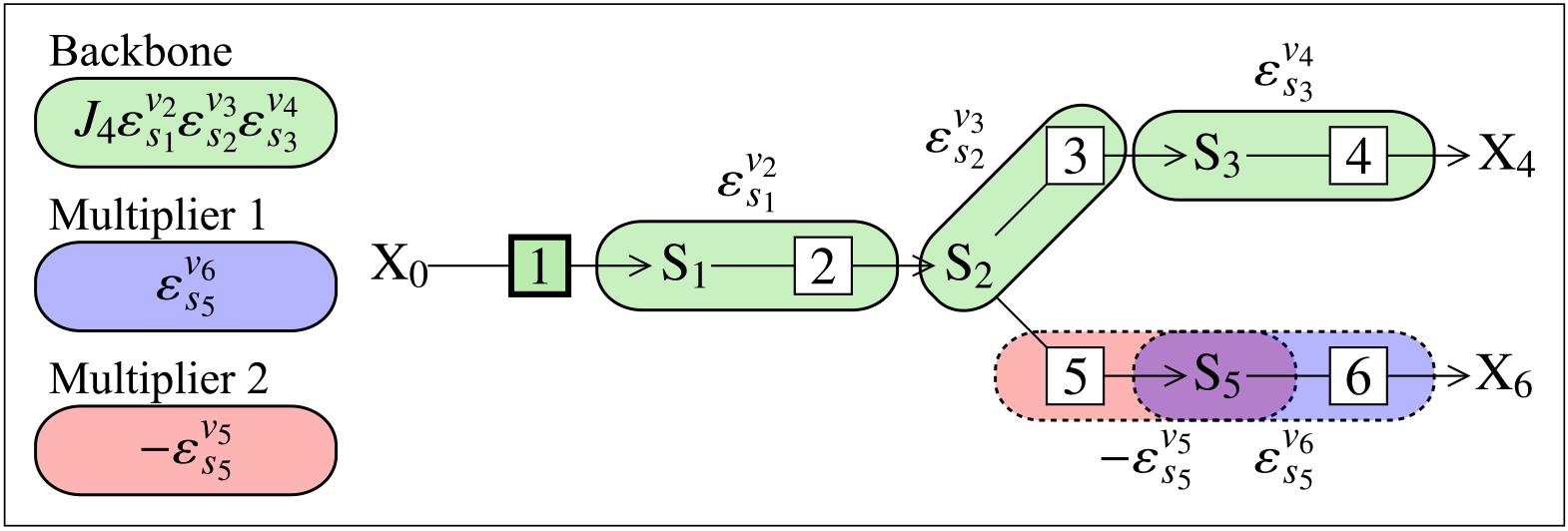
Control patterns for a 6-step branched metabolic pathway. Backbone and multiplier patterns for two control patterns of 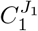 are depicted. Green bubbles indicate the backbone pattern, while red and blue bubbles each indicate a different multiplier pattern. Each multiplier-backbone combination (Backbone × Multiplier 1 and Backbone × Multiplier 2) represents a single control pattern. The chain of local effects for the backbone originates from a perturbation of *v*_*1*_, which, if positive, causes an increase in *s*_1_, which increases *v*_2_, then *s*_2_, then *v*_3_, then *s*_3_, and, finally, *v*_4_; each of these effects plays a role in determining the sensitivity of *J*_1_ towards *v*_1_ through the backbone, which in turn is modified by one of the two multiplier patterns.

In this paper we consider the absolute values of the control patterns for determining their contribution to the associated control coefficient. This contribution is calculated as the percentage of the absolute value of the control pattern as a fraction of the sum of the absolute values of all control patterns for that control coefficient. This metric is used instead of a conventional percentage to account for cases where control patterns have different signs, which could lead to the contribution of a single pattern being more than 100%.

Control patterns of branched pathways can be factored into subpatterns called the backbone and multiplier patterns [38]. In the case of flux control coefficients, a backbone pattern is defined as an uninterrupted chain of reactions that links two terminal metabolites and passes through the flux being controlled (the reference flux). Multiplier patterns are chains of reactions that occur in branches to a backbone pattern, and so occur only in branched pathways. One backbone pattern can be combined with various multipliers to form different control patterns, and a single multiplier can be associated with control patterns with different backbones.

### Regulation by enzymes

One definition of regulation, as it pertains to metabolic systems, is *the alteration of reaction properties to augment or counteract the mass-action trend in a network of reactions* [17]. Enzyme activity represents one of the means by which the mass-action trend can be counteracted, with higher potential for regulation being achieved far from equilibrium. Thus, it is necessary to be able to determine a reaction’s distance from equilibrium, and to be able to distinguish between the effect of mass action and enzyme binding in order to quantify the regulatory effect of enzymes in a system [16].

Distance from equilibrium is given by the disequilibrium ratio *ρ* = Г/*K*_*eq*_, where Г is the mass-action ratio (also termed reaction quotient). Kinetic control in the forward direction is indicated by *ρ* ≤ 0.1, thermodynamic control in the forward direction is indicated by *ρ* ≥ 0.9, and a combination of kinetic and thermodynamic control is indicated by 0.1 < *ρ* < 0.9 [16].

Recasting a rate equation into logarithmic form allows one to separate the effects of binding and mass action on the reaction rate into two additive terms. Partial differentiation of the logarithmic rate equation with respect to a substrate or a product yields two elasticity coefficients, one of which quantifies the effect on reaction rate of binding of substrate or product by the enzyme, and the other the effect of mass action [16,17]. A substrate elasticity, for example, will be partitioned as

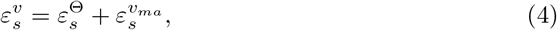

where 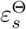 represents the binding elasticity and 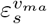 the mass-action elasticity components [16]. Similarly, for a product, 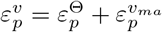.

### Software

Model simulations were performed with the Python Simulator for Cellular Systems (PySCeS) [39] within a Jupyter notebook [40]. Symbolic inversion of the **E** matrix (Eq. 3) and subsequent identification and quantification of control patterns was performed by the SymCA [12] module of the PySCeSToolbox [13] add-on package for PySCeS. Control patterns for each control coefficient were automatically numbered by SymCA starting from 001, and we used these assigned numbers in the presented results. Importantly, SymCA was set to automatically replace zero-value elasticity coefficients with zeros. In other words, certain elasticity coefficients are never found within any control coefficient expressions as their value will always be zero (such as in the case of elasticity coefficients of irreversible reactions with respect to their products).

Additional manipulation of symbolic expressions, data analysis, and visualisations were performed using the SymPy [41], NumPy [42] and Matplotlib [43] libraries for Python [44].

### Model

As mentioned, results obtained during a previous study of a model of pyruvate branch metabolism in *Lactococcus lactis* [29] were revisited in this paper. This model was originally constructed by Hoefnagel *et al.* [21], and was obtained from the JWS online model database [45] in the PySCeS model descriptor language (see http://pysces.sourceforge.net/docs/userguide.html). A scheme of the pathway is shown in Fig. 2.

Members of the of ATP/ADP, acetyl-CoA/CoA and NADH/NAD^+^ moiety-conserved cycles were treated as ratios in order to perform parameter scans of these conserved moieties without breaking moiety conservation. The notation *ϕ*_*A*_, *ϕ*_*C*_, and *ϕ*_*N*_ (see Fig. 2) will be used henceforth. Here, we only considered the effect of changing *ϕ*_*N*_. The value of *ϕ*_*N*_ was thus fixed and varied between 2 × 10^−4^ and 1.77 in order to generate the results presented in this paper.

The value of *ϕ*_*N*_ was varied directly rather than modulating its demand (the activity of NADH oxidase) for two reasons. First, we wanted to mirror the methodology used in our original study [29], so that we could further explore the results obtained there. Second, we wanted to simplify the system for control-pattern analysis. These considerations are discussed further at the end of the Results section, where we perform a similar control-pattern analysis for a range of *ϕ*_*N*_ values by varying the *V*_*max*_ of NADH oxidase in a version of the model where *ϕ*_*N*_ is not fixed.

**Fig 2.**
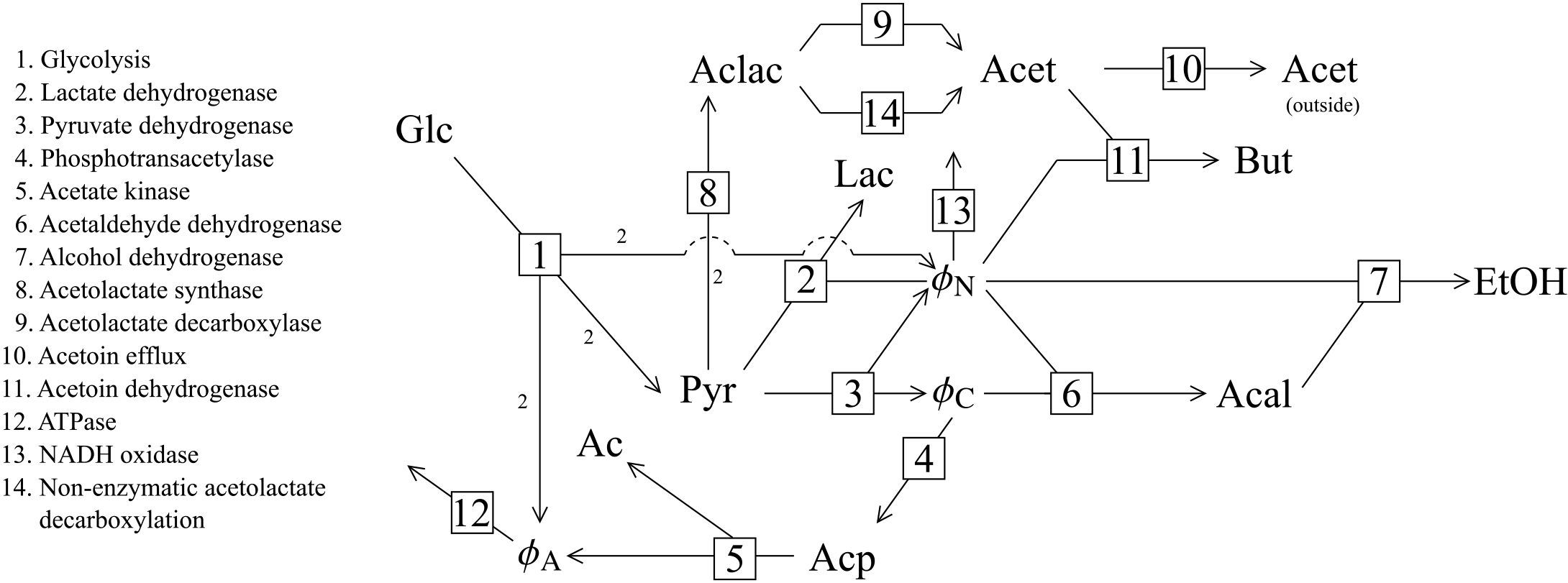
The pyruvate branch pathway. Reactions are numbered according to the key. For the sake of brevity we refer to the reactions by their number instead of their name throughout this paper. The stoichiometry of each reaction is 1 to 1, except for reaction 1 where Glc + 2ADP + 2NAD^+^ → 2Pyr + 2ATP + 2NADH and reaction 8 where 2Pyr ⇌ Aclac. Intermediates are abbreviated as follows: Ac: acetate; Acal: acetaldehyde; Acet: acetoin; Aclac: acetolactate; Acp: acetyl phosphate; Glc: glucose; Lac: lactate; But: 2,3-butanediol; Pyr: pyruvate; EtOH: ethanol; *ϕ*_*A*_ ATP/ADP; *ϕ*_*C*_: acetyl-CoA/CoA; *ϕ*_*N*_: NADH/NAD^+^.

## Results

A main finding of our previous study [29] was that an increase in *ϕ*_*N*_ caused a decrease in the flux through the acetaldehyde dehydrogenase reaction block (*J*_6_) in spite of NADH being a substrate for reaction 6. Investigation of the partial response coefficients of *J*_6_ with respect to *ϕ*_*N*_ revealed this unintuitive effect to be the result of the interaction of *ϕ*_*N*_ with pyruvate dehydrogenase (reaction 3) as signified by the large negative partial response coefficient 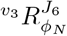. This route of interaction was found to be one of the most dominant effects in the regulation of *J*_6_ by *ϕ*_*N*_.

Dividing the dominant partial response coefficients into their component control and elasticity coefficients illustrated their contributions to the overall observed response. In simple terms we could now understand that the negative *J*_6_ response was due to the rate of reaction 3 responding negatively towards a decrease in its substrate concentration (NAD^+^) which, together with the relatively large flux control of reaction 3 on *J*_6_ 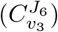, caused the overall negative *J*_6_ flux response.

In the following sections we will continue this line of investigation by examining how 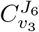 is determined by the interactions between the various species and enzymes of the system.

### Identification of dominant control patterns of 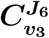

Using the SymCA software tool, we identified 76 control patterns for 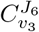 and generated expressions for each. While this is a much smaller number than the 226 patterns identified in the system where *ϕ*_*N*_ was a free variable, a naive investigation into the properties of each would still represent an unwieldy task. We therefore selected for further investigation only the most important control patterns in terms of their contribution towards 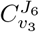 for the tested range of *ϕ*_*N*_ values.

Two cut-off values were used to select the most important control patterns: not only did such patterns have to exceed a set minimum percentage contribution towards the sum of the control patterns, they also had to exceed this first cut-off value over a slice of the complete *ϕ*_*N*_-range; the second cut-off set the magnitude of this slice as a percentage of the *ϕ*_*N*_-range on a logarithmic scale. The first cut-off excludes control patterns that make a negligible contribution to the sum of control patterns, while the second ensures that those that make the first cut do so over a slice of the *ϕ*_*N*_-range. Selection of the two cut-off values was automated by independently varying their values between 1% and 15% in 1% increments and selecting the smallest group of control patterns that could account for at least 70% of the absolute sum of all the control patterns. While these criteria are admittedly somewhat arbitrary, they reduced the number of control patterns to consider in a relatively unbiased manner while still accounting for the majority of control.

The two cut-off values of 7% and 5% yielded 11 important control patterns, the smallest group of all the different cut-off combinations. These patterns, as shown in Fig. 3B, made varying contributions towards 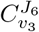 depending on the value of *ϕ*_*N*_, with different patterns being dominant within different ranges of *ϕ*_*N*_ values. The control patterns were categorised into groups numbered 1-4 based on the range of *ϕ*_*N*_ values in which they were responsible for the bulk of the control coefficient value, with each group thus dominating the overall value of 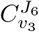 within their active range. Patterns in group 3 (CP063 and CP030) were an exception to this observation as they shared dominance with groups 2 and 4 in their active range. Fig. 3A provides another perspective on the contribution of the 11 dominant control patterns towards the control coefficient by demonstrating how closely their sum approximates the value of 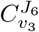.

While it is clear that, depending on the value of *ϕ*_*N*_, certain control patterns are more important than others, and that they can be grouped according to similar responses towards *ϕ*_*N*_, it is impossible to understand how this behaviour arises without investigating their actual composition and structure. In the next section we will address this issue and begin to dissect the control patterns based on the contributions of their individual components.

### Backbone and Multiplier Patterns of 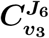

To investigate the source of the differences between the 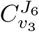 control patterns we subdivided them into their constituent backbone and multiplier patterns, since these subpatterns provide an intermediate level of abstraction between the full control patterns and their lowest level flux and enzyme elasticity components. This process yielded six backbone patterns and 13 multiplier patterns as shown in Table 1. Multiplier patterns were categorised into two groups labelled “T” and “B” based their components being located in the top or the bottom half of the pathway scheme in Fig. 2. Each of the 76 control patterns consisted of a single backbone pattern and either one (B) or two multiplier (B and T) patterns as can be seen in Table 2.

Control patterns within each of the four dominant control pattern groups (as indicated in Fig. 3) were found to be related in terms of their subpattern composition (see Table 2): each control pattern within the same group had the same backbone and T multiplier, except for control patterns in group 2, which did not contain any T multipliers and thus only shared a backbone pattern. The backbone pattern of group 2 (B in Table 1) did, however, extend into the same metabolic branch as the T multipliers via acetolactate synthase (reaction 8), thus effectively acting as both a backbone and T multiplier for this group of control patterns. Control patterns within each group therefore differed only in terms of their B multipliers. Furthermore, T multipliers were unique to each group and only groups 3 and 4 shared the same backbone. On the other hand, the same B multipliers were found in the control patterns of multiple groups, with B3 forming part of the dominant pattern in each group. As would be expected from the low number of control patterns making up the bulk of the observed control, a large number of backbone and multiplier patterns do not appear in any of the important control patterns.

**Fig 3.**
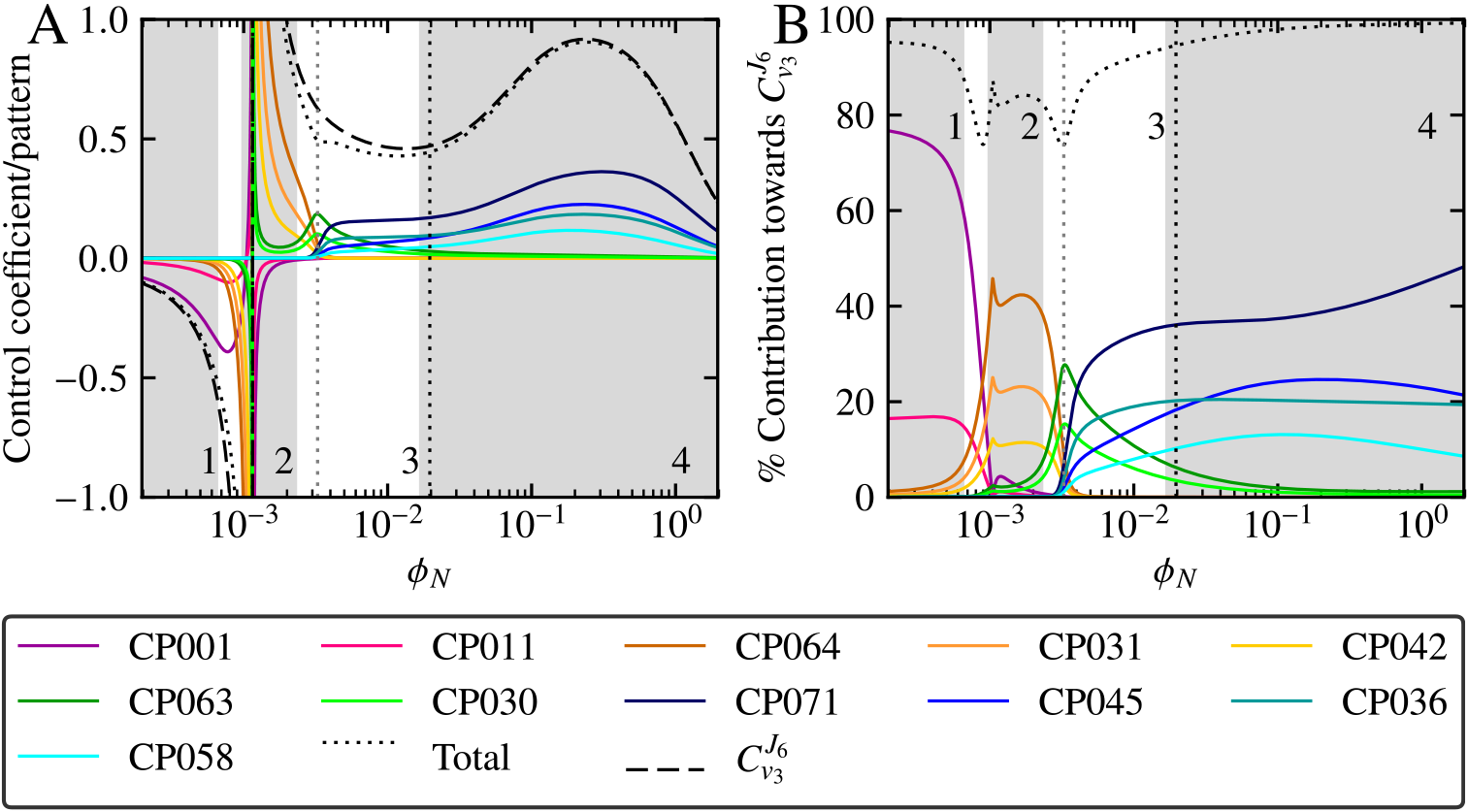
The most important control patterns of 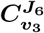 as functions of *ϕ*_*N*_. Control patterns were chosen according to the criteria described in the text. (A) The most important control patterns shown in relation to the value of 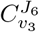 and the value of their total sum. (B) The absolute percentage contribution of the most important control patterns relative to the absolute sum of the values of all 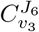 control patterns. Control patterns and 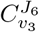 are indicated in the key. In both (A) and (B) grey shaded regions indicate ranges of *ϕ*_*N*_-values that are dominated by a group of related control patterns, while unshaded regions indicate shared contribution by unrelated patterns. Groups are numbered 1-4 (see Table 1), and control patterns belonging to a group are colour coded such that group 1 is pink, 2 is yellow, 3 is green, and 4 is blue. Control in the unshaded region 3 is shared between the group 2 and 3 patterns for *ϕ*_*N*_ < 0.0033 and by the group 3 and 4 patterns for *ϕ*_*N*_ > 0.0033 as indicated by the vertical dotted gray line. The switch from negative control coefficient values to positive values indicates indicates the reversal of direction of *J*_6_ flux. The black dotted vertical line indicates the steady-state value of *ϕ*_*N*_ in the reference model.

A parameter scan of *ϕ*_*N*_, shown in Fig. 4, highlighted two important features of the backbone and multiplier patterns. First, some patterns differed vastly in terms of magnitude when compared to members of their own type of pattern as well as when compared to other types of patterns. The up to four orders of magnitude of difference between B3 and B6 (Fig. 4B) is illustrative of the former case, whereas the ~ 20 orders of magnitude difference between B3 and backbone E (Fig. 4B and A) illustrates the latter. Second, all patterns within a group tended to respond similarly towards increasing *ϕ*_*N*_, with the B multipliers remaining relatively constant (Fig. 4B) and the T multipliers deceasing in magnitude over the tested *ϕ*_*N*_-value range (Fig. 4C). The common-denominator-scaled backbone patterns were an exception to this second observation since two (A and C) increased in magnitude in response to increasing *ϕ*_*N*_ while the rest (B, D, E and F) decreased in magnitude (Fig. 4A). However, the unscaled backbone patterns all decreased in magnitude for the *ϕ*_*N*_ range. It is important to note that while scaling by the common denominator is necessary to account for the all the components of the control pattern expressions, the choice to scale the backbone patterns (as opposed to either the T or B multipliers) was made arbitrarily. The magnitudes of the backbone patterns relative to each other are unaffected by scaling, and general observations regarding the responses of the scaled backbone patterns towards changes in *ϕ*_*N*_ or alterations in the system hold for the unscaled backbone patterns.

**Table 1.**
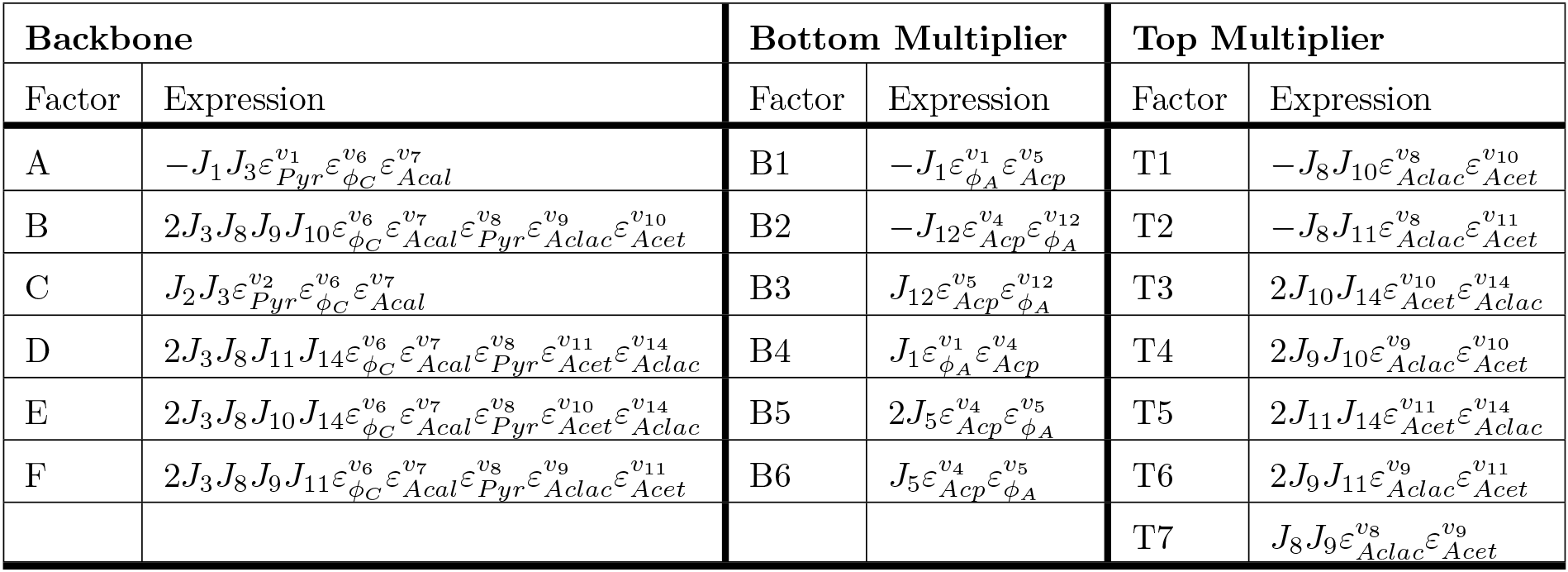
Backbone and multiplier expressions of the control patterns of 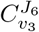. Backbone expressions are named A-F and multipliers are classified according to their relative position on the reaction scheme, with B and T multipliers appearing on the bottom and top halves respectively. Each of the control pattern numerators of 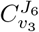 is a product of one of the backbones and either a B (for backbones B, D-F) or both a B and a T multiplier (for backbones A and C). Valid control pattern numerators do not consist of multiplier-backbone combinations with overlapping factors.

An explicit example of how the backbone and multiplier patterns determined the values of control patterns is displayed in Fig 4D, which shows CP063 and CP071 together with their subpattern components as a function of *ϕ*_*N*_. Here it is important to note that backbone and multipliers with values larger than 1 act to increase the value of their control pattern, while those with values below 1 act to decrease the control pattern value. Since each order of magnitude above or below 1 has the same relative positive or negative effect on the control pattern, the use of a logarithmic scale in Fig 4 allows us to easily gauge the effects of the subpatterns on the overall control pattern magnitude. While CP063 and CP071 both made large contributions towards 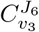 in region 3, as indicated in Fig 3B, CP071 had a much larger value than CP063 for most *ϕ*_*N*_ values in spite of only differing in terms of their T multiplier components (with CP071 containing T6 and CP063 containing T4). While the steady increase in the magnitude of backbone C clearly played a role in the dominance of these two control patterns within their respective *ϕ*_*N*_ ranges, we can see that the larger T4 value at *ϕ*_*N*_≈ 3 × 10^−3^ caused CP063 to dominate at this point, while the decrease of T4 to a lower magnitude than T6 for *ϕ*_*N*_ values larger than this point caused CP071 to dominate for the remainder of the series of *ϕ*_*N*_ values. In essence, T4 pulled the value of CP063 down more than T6 did CP073 in region 4. The matching hatched regions between CP063 and CP071, and T4 and T6, clearly show that the difference between the two control patterns can be attributed solely to these two T multipliers.

A more complicated example of the same principle can be seen in Fig 4E and F which respectively shows the control patterns CP001 and CP071, and their constituent subpatterns. Unlike CP063 and CP071 (Fig 4D), which only differed in terms of a single subpattern, CP001 and CP071 differed in terms of both their backbone and their T multiplier components, with CP001 consisting of T1, A and B3, and CP073 consisting of T6, C and B3. The dominance of CP001 compared to CP071 in region 1 of Fig 3 can now be seen to be a result of the large values of T1 and A in this *ϕ*_*N*_ range compared to their counterparts, T6 and C. On the other hand, within the *ϕ*_*N*_ range represented by regions 3 and 4 in Fig 3, both T6 and C had larger values than T1 and A, therefore causing the dominance of CP071 over CP001 within these two regions. Similar to Fig 4D, the difference in magnitude between T1 and T6, and between A and C are shown as diagonal hatching in Fig 4F, with the combined effect of these differences being shown as cross hatching in Fig 4F, thus illustrating how the differences between these subpatterns contribute to the overall difference in magnitude between CP001 and CP071.

**Fig 4.**
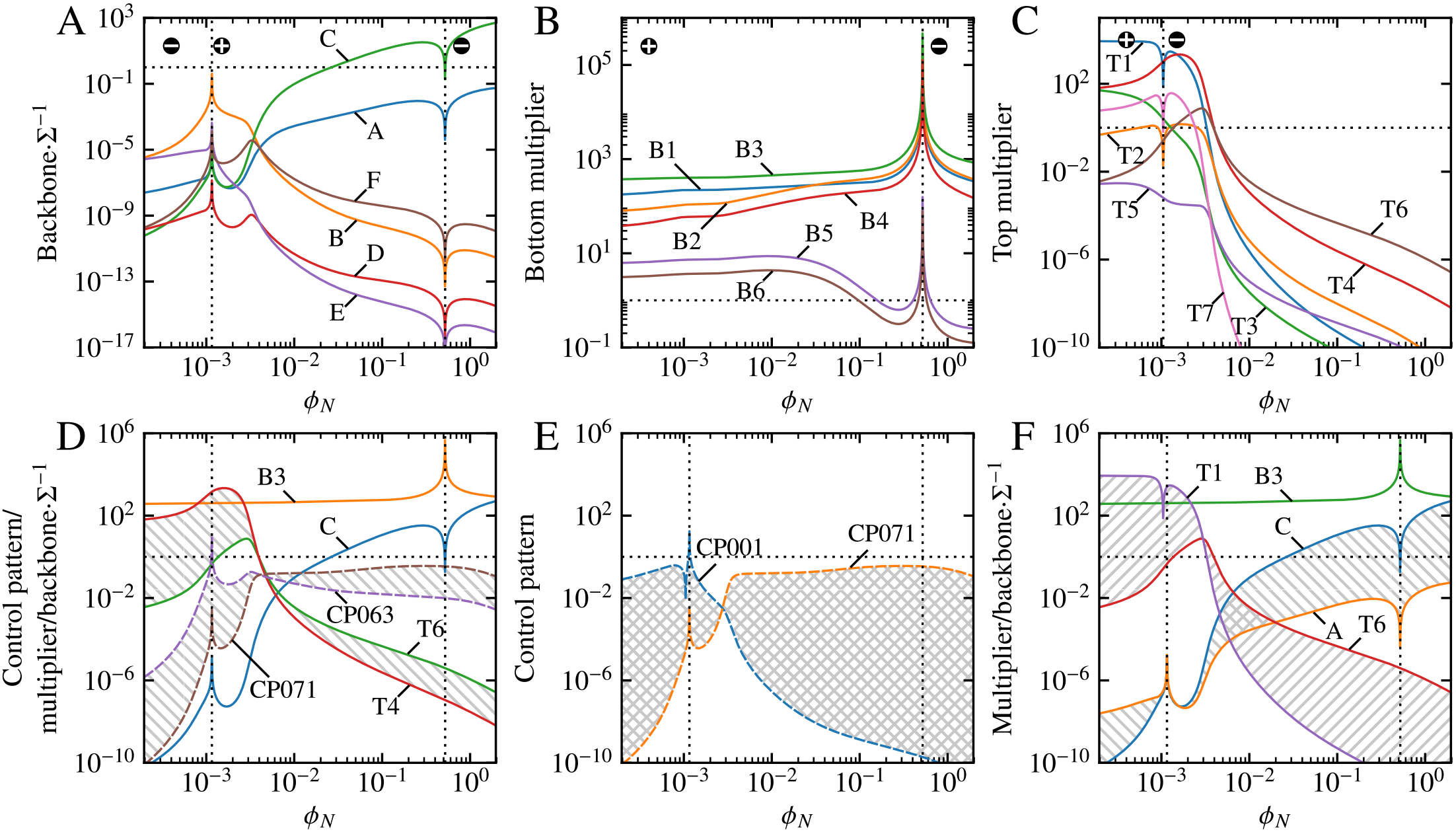
Backbone and multiplier patterns of the 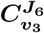 control patterns as functions of *ϕ*_*N*_. (A) Values of the backbone patterns (A-F) scaled by the control coefficient common denominator (Σ) of this pathway. (B) Values of the multiplier patterns consisting of components from the bottom half of the reaction scheme in Fig. 2 (B1-B6). (C) Values of the multiplier patterns consisting of components from the top half of the reaction scheme (T1-T7). (D) CP063 and CP071 together with their constituent scaled backbone and multiplier patterns. CP063 and CP071 differ only in their top multipliers (T4 and T6), as is reflected by the corresponding hatched areas between the pairs CP063/CP071 and T4/T6. (E) CP001 and CP071. (F) The constituent scaled backbone and multiplier components of CP001 and CP071. CP001 and CP071 differ both in terms of their backbones (A and C) and their top multipliers (T1 and T6); these differences are indicated with hatching between T1/T6 and A/C, and their cumulative effect is indicated with hatching between CP001/CP071 in (E). The horizontal black dotted line at *y* =1 differentiates between patterns that have an increasing (*y* > 1) or diminishing (*y* < 1) effect on their control pattern products. Absolute values of pattens are taken to allow for plotting logarithmic coordinates; the crossover from positive to negative values with increasing *ϕ*_*N*_ is indicated by vertical dotted lines. Backbone patterns each switch sign twice from a negative starting point on the left-hand side of (A). The multiplier patterns T3, T4, T5, and T6 are positive throughout the *ϕ*_*N*_ range, whereas the remaining T multipliers switch from positive to negative.

**Table 2.** Numerator expressions of the dominant control patterns of 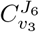. The numerators of the control patterns highlighted in Fig. 3 are expressed in terms of their constituent backbone and multiplier factors (Table 1) and are separated into groups based on the *ϕ*_*N*_ ranges for which they are most important, as in Fig. 3. Control patterns are arranged in descending order of relative importance within their group.

| Group | Control Pattern | Expression |
| --- | --- | --- |
| 1 | CP001 | $A \cdot B3 \cdot T1$ |
| | CP011 | $A \cdot B2 \cdot T1$ |
| 2 | CP064 | $B \cdot B3$ |
| | CP031 | $B \cdot B1$ |
| | CP042 | $B \cdot B2$ |
| 3 | CP063 | $C \cdot B3 \cdot T4$ |
| | CP030 | $C \cdot B1 \cdot T4$ |
| 4 | CP071 | $C \cdot B3 \cdot T6$ |
| | CP045 | $C \cdot B2 \cdot T6$ |
| | CP036 | $C \cdot B1 \cdot T6$ |
| | CP058 | $C \cdot B4 \cdot T6$ |

These results show how the combined effect of different subunits of a metabolic pathway (i.e., those represented by backbone and multiplier patterns) determines the control patterns of a given control coefficient. In our case, the subpatterns clearly correspond to different metabolic branches and thus narrow down the search for the ultimate source of the differences in behaviour between different control patterns.

### Following the chains of effects

In this section we relate the control patterns discussed above to their constituent elasticity coefficients, thus demonstrating how the properties of the most basic components of a metabolic system (i.e. enzyme-catalysed reactions) determine the systemic control properties as quantified by control coefficients.

As previously mentioned, CP071 and CP063 differ only in terms of their T multiplier patterns, T6 and T4. However, upon closer inspection of the composition of these two multiplier patterns (Table 1) it became clear that these patterns were very similar, with both containing a 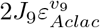 factor. The only difference is that T6 had 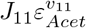 as a factor, whereas T4 had 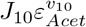. A visual representation of the components of CP071 and CP063, as shown in Fig. 5, serves to further illustrate how closely related these two control patterns are.

Fig. 6A shows the individual components of the T4 and T6 as a function of *ϕ*_*N*_. For much of the *ϕ*_*N*_ range below approximately 4 × 10^−3^, both the T4-exclusive factors *J*_10_ and 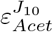 were multiple orders of magnitude larger than their T6 counterparts. This clearly accounted for the dominance of CP063 over CP071 in this *ϕ*_*N*_ range, and indeed for the dominance of the whole of group 3 over group 4 in this range, since all the patterns in each of these two groups shared the T4 multiplier. As *ϕ*_*N*_ increased, however, the two fluxes *J*_10_ and *J*_11_ become practically equal both in magnitude and in terms of their responses towards *ϕ*_*N*_ (except for *ϕ*_*N*_ values around 8.5 × 10^−3^ where *J*_11_ switched from positive to negative). This means that the observed regime change from CP063 to CP071, and from the group 3 to group 4 patterns, with increasing *ϕ*_*N*_ was completely determined by the difference in sensitivity of reactions 10 (acetoin efflux) and 11 (acetoin dehydrogenase) towards their substrate as represented by 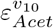 and 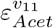 respectively.

**Fig 5.**
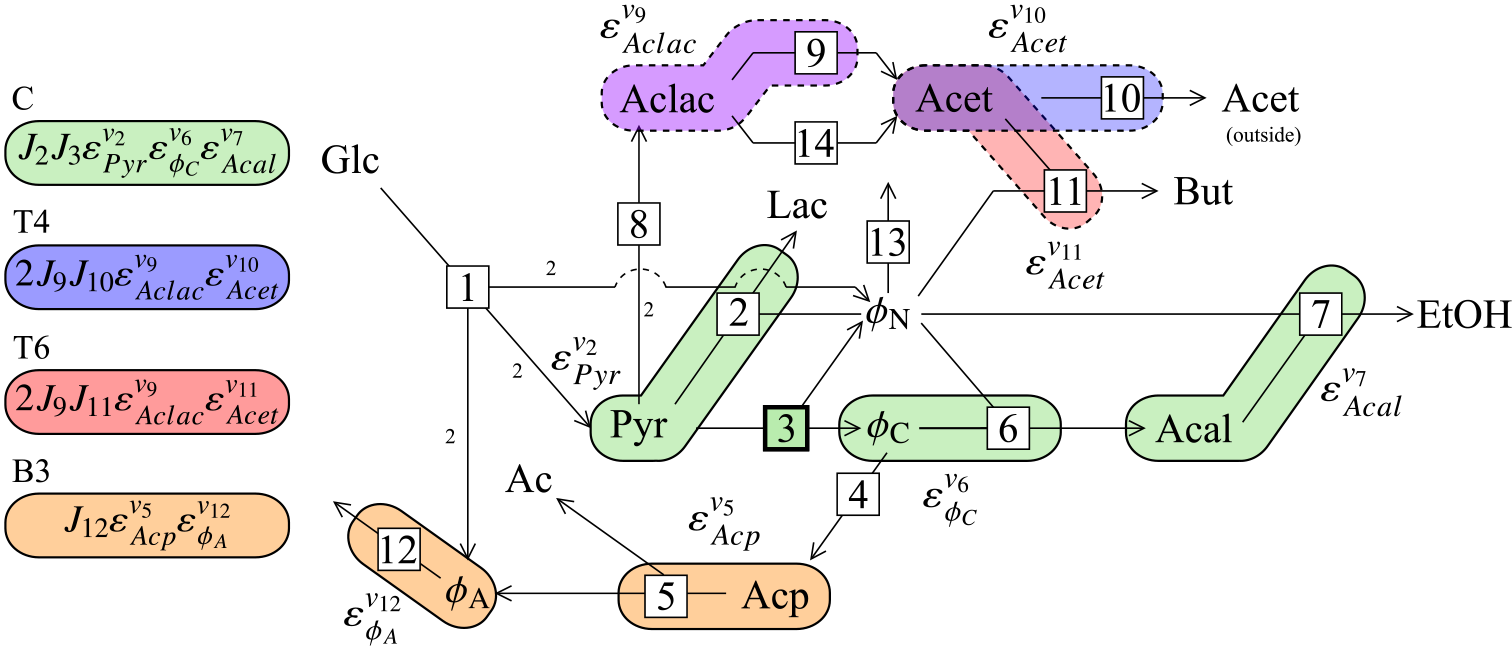
The components of control patterns 063 and 071 of 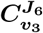. Backbone and multiplier patterns that constitute the dominant control patterns of group 3 and group 4 (CP063 and CP071) are indicated as groups of coloured bubbles that highlight their elasticity coefficient components as indicated by the key. These control patterns both share backbone C and multiplier B3 and are therefore differentiated based on their incorporation of either multiplier T4 (CP063) or T6 (CP071).

The observed differences between 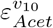 and 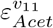 can be explained by examining the properties of reactions 10 and 11 within the thermodynamic/kinetic analysis framework [16]. The value of 1 of 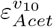 for *ϕ*_*N*_ ≳ 4 × 10^−3^ was the result of two factors: first, the concentration of the substrate for reaction 10 (Acet) was far from saturating within this range 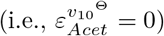, and second, reaction 10 was defined as an irreversible reaction in this system, i.e., 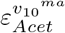 always has a value of 1. Thus 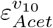 was determined by both enzyme binding and the mass-action effect for *ϕ*_*N*_ ≲ 4 × 10^−3^, and exclusively by mass-action for *ϕ*_*N*_ ≳ 4 × 10^−3^.

Reaction 11 was defined as a reversible bi-bi reaction, meaning that 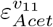 could potentially exhibit more complex behaviour than its counterpart. From the 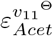 value of −1 for *ϕ*_*N*_ ≲ 3 × 10^−3^ (Fig. 6B), it is clear that the concentration of Acet remained fully saturating for reaction 11 within this range. Because *ρ* had a value much smaller than 1 for this *ϕ*_*N*_ range, 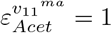. As *ϕ*_*N*_ increased, the value of 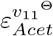 increased to zero, indicating a decrease in the effect of substrate binding, which ultimately led to a zero contribution towards the overall value of 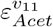 for *ϕ*_*N*_ ≳ 5 × 10^−3^. Similarly, 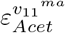 also increased in response to increasing *ϕ*_*N*_, tending towards ∞ as equilibrium was approached (*ρ* = 1). A further increase in *ϕ*_*N*_ beyond the point of equilibrium caused 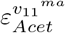 to become negative with a relatively large magnitude, indicating that reaction 11 remained close to equilibrium for the remaining *ϕ*_*N*_ values.

**Fig 6.**
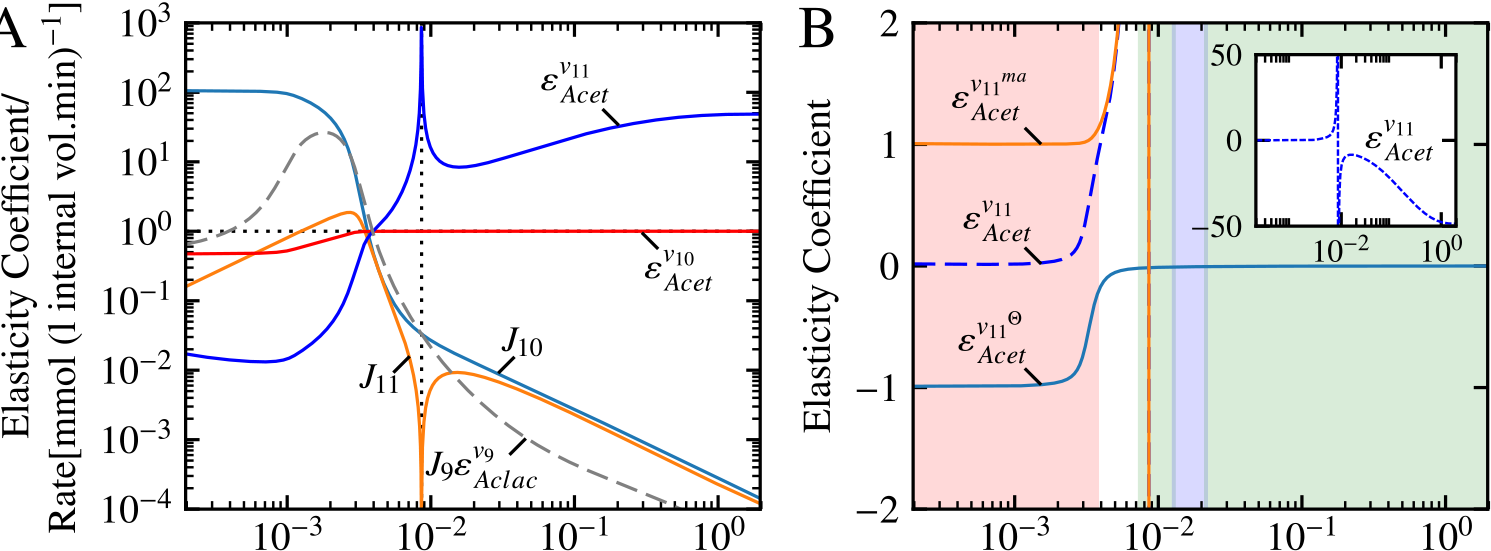
The flux and elasticity components of T4 and T6 as functions of *ϕ*_*N*_. (A) The elasticity and flux factors that constitute the multipliers T4 and T6. Both multipliers share the factor 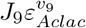, whereas *J*_10_ and 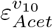 are unique to T4, and *J*_11_ and 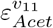 are unique to T6. Logarithmic coordinates together with a horizontal black dotted line and absolute values are used in the same fashion as in Fig. 4 with the black vertical dotted line indicating the crossover from positive to negative values of *J*_11_ and 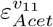. (B) The elasticity coefficient 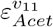 split into its binding and mass action components. The insert shows 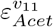 on an expanded scale. Shaded areas indicate kinetic vs. thermodynamic control of *υ*_11_ with red: *ρ* ≤ 0.1, white: 0.1 < *ρ* < 0.9, green: 0. 9 ≤ *ρ* ≤ 1/0.9 and blue: 1/0.9 < *ρ* < 1/0.1.

As a second example we will revisit the differences between CP001 and CP071 as discussed in the previous section. There are many more differences between these two control patterns than between CP063 and CP071, since both the backbone and T multiplier patterns are responsible for determining the regions for which they are dominant. The differences between CP001 and CP071 are visually depicted in Fig. 7. Figs. 8A and B shows the individual components of the backbone and T multiplier respectively belonging to CP001 and CP071. Note that the components *J*_3_, 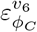, and 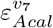 appears as components in both backbone A and C and are thus collected into a single factors. For the sake of clarity we will refer to factors that act to increase the magnitude of a control pattern as “additive components” and those that act to decrease its magnitude as “subtractive components”, since these terms reflect the effect of the factors in logarithmic space. If we focus on the left-hand side of Figs. 8A and B where CP001 dominated, it is clear that this control pattern has more additive components (*J*_1_, *J*_8_, and *J*_i0_) than CP071 (*J*_2_, and *J*_9_). Additionally, for these low *ϕ*_*N*_ values, the subtractive components of CP001 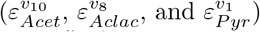 had values closer to 1 than those of CP071 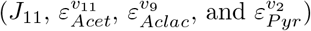, thus having a smaller diminishing effect on CP001. As *ϕ*_*N*_ increased, however the situation reversed. Focusing on the region where *ϕ*_*N*_ > 3 × 10^−3^ and CP071 is larger than CP001, we see that all but one (*J*_1_) of the components that were additive for lower *ϕ*_*N*_ values decreased by more than 6 orders of magnitude, thus becoming subtractive. Similarly, while the value of 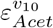 increased to ≈ 1, the remaining subtractive components greatly decreased in magnitude for this range. On the other hand, while one of CP071’s subtractive components (*J*_11_) and one of its additive components (*J*_9_) both decreased in value (with *J*_9_ becoming subtractive), all other components increased in value. Therefore for the region where CP071 dominated it had two large additive components and two subtractive components compared to CP001’s single additive and four subtractive components.

These changes in the values of the subtractive and additive components of CP001 and CP071 as *ϕ*_*N*_ increased can largely be explained in terms of two changes in flux. First, flux was diverted from the acetoin producing branch (*J*_8_) towards the lactate producing branch as *ϕ*_*N*_ increased due to *ϕ*_*N*_ acting as a co-substrate for *v*_2_. Second, *J*_1_ decreased due to product inhibition of *υ*_1_ by *ϕ*_*N*_, resulting in a much larger decrease in *J*_8_ relative to *J*_2_. This shift in flux also caused decreases in *J*_8_ and *J*_10_ (belonging to CP001), and *J*_9_ and *J*_11_ (belonging to CP071). These changes in flux also had effects on the elasticity coefficients. As previously discussed, 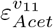 increased due to *υ*_11_ becoming close to equilibrium. The elasticity coefficient 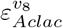 decreased in magnitude due to a decrease in Aclac concentration as a result of decreased *J*_8_. The decrease in Aclac also caused 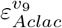 to increase to a value of one. Similarly, the decrease in Pyr partly due to decreased *J*_1_ resulted in the decrease in 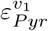 and the increase in 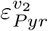. Thus, broadly speaking, the shift in dominance of control patterns in group 1 to those in group 4 can be attributed to the shifts in flux between the pyruvate-consuming and -producing branches of the system.

**Fig 7.**
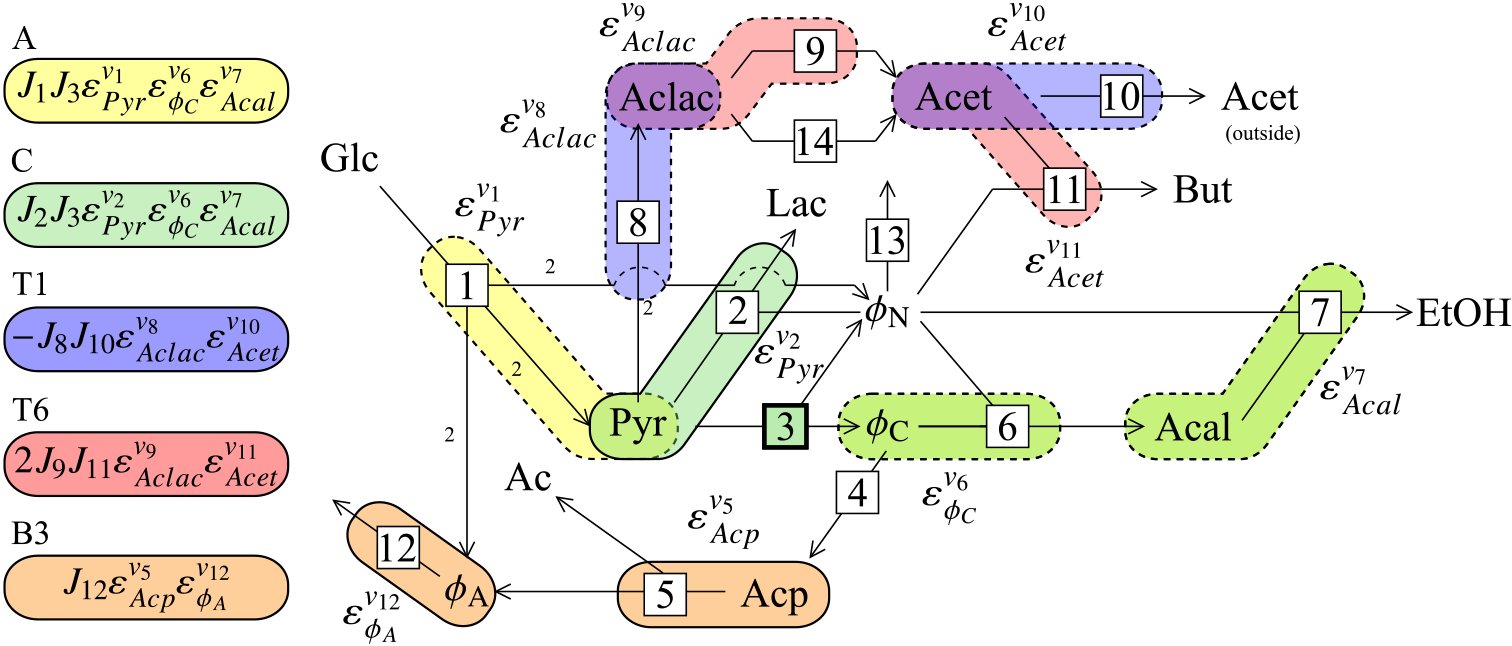
The components of control patterns 001 and 071 of 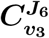. Backbone and multiplier patterns that constitute the dominant control patterns of group 1 and group 4 (CP001 and CP071) are indicated as groups of coloured bubbles that highlight their elasticity coefficient components as indicated by the key. These control patterns have different backbone patterns (A and C), as well as different multipliers (T1 and T6).

**Fig 8.**
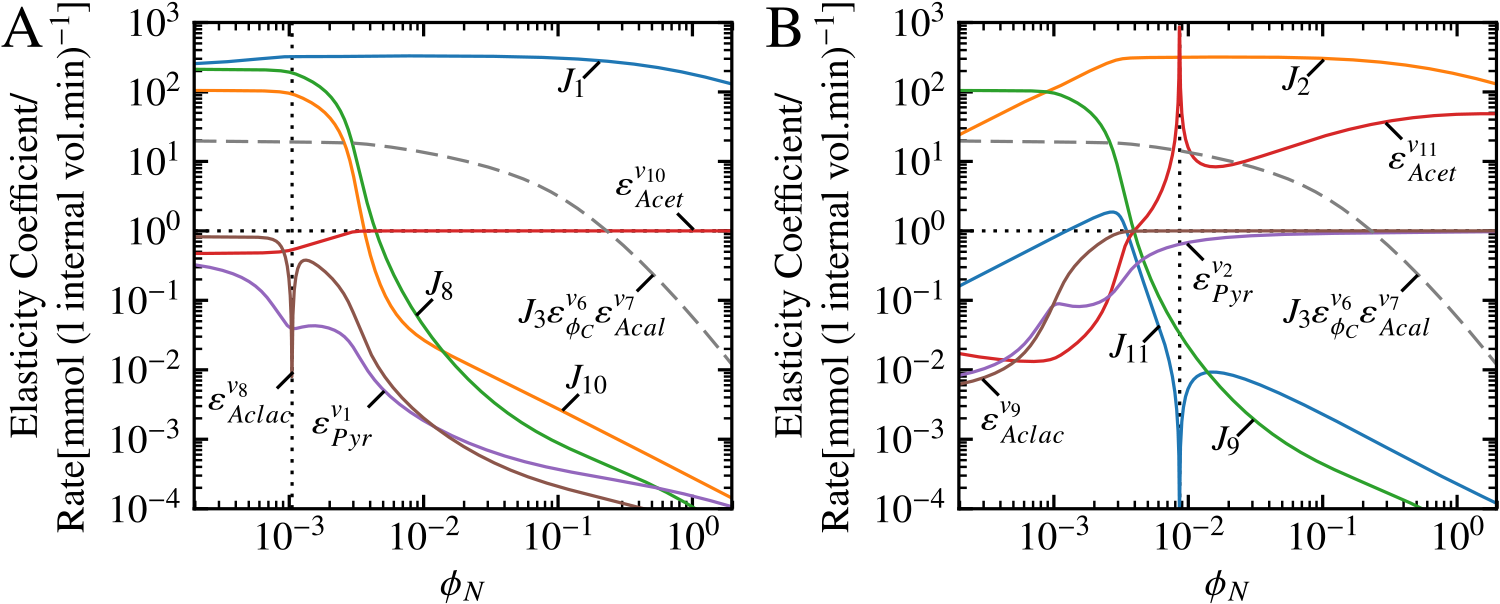
The flux and elasticity components of T1 × A and T6 × C as functions of *ϕ*_*N*_. (A) The elasticity and flux factors that constitute the multiplier T1 and backbone A. (B) The elasticity and flux factors that constitute the multiplier T6 and backbone C. Both T1 × A and T6 × C share the factor 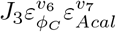, which is indicated as a single dashed grey line in each of (A) and (B). Logarithmic coordinates together with a horizontal black dotted line and absolute values are used in the same fashion as in Fig. 4. The the black vertical dotted line in (A) indicates crossover from positive to negative values of 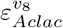 (due to product activation as a result of cooperative product binding in the reversible Hill equation [46]) and in (B) indicates the crossover from positive to negative values of *J*_11_ and 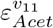.

### Altering control via manipulation of system components

The results above demonstrate how the properties of single reactions in a metabolic pathway contribute to the behaviour of a system on a global scale. In the final section we will use this information to demonstrate and assess a possible strategy for altering the control properties of a metabolic system.

Since the difference between the factors 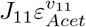 and 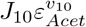 was the source of the difference in contribution of CP071 and CP063 towards 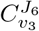, we can imagine that altering these factors would have an impact on the control of the system. If we were to knock out acetoin dehydrogenase and replace reaction 11 with an hypothetical enzyme catalysing a reaction with a thousand-fold larger *K*_*eq*_ value, we would expect this new reaction to be far from equilibrium for the same *ϕ*_*N*_ values where our original reaction was near equilibrium. While such an alteration in *K*_*eq*_ is unrealistic, it is reasonable to expect that such a change could affect the values of *J*_11_ and 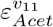, ultimately leading to a change in the control properties of the system.

In reality, however, the situation is not so simple. Changing the value of *K*_*eq*_ of reaction 11 from 1.4 × 10^3^ to 1.4 × 10^6^ did indeed yield some positive results. First, 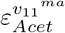 had a value of one for almost the complete range (S1B Fig.), which led to 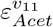 having practically the same value as 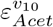 for *ϕ*_*N*_ ≳ 4 × 10^−3^. Second, *J*_11_ had a lower magnitude than previously for *ϕ*_*N*_ ≳ 4 × 10^−3^ and the flux did not reverse (S1A Fig.). However, a number of unintended side effects also occurred. Most notably *J*_10_ decreased so that its value was lower than that of *J*_11_, thereby offsetting the decrease in *J*_11_ and 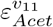. Thus, while these changes slightly lowered the values of both T6 and T4, their values relative to each other remained almost unchanged (S3C Fig.). Additionally, the backbone pattern C exhibited an increase in magnitude as a result of the new *K*_*eq*_ value (S3A Fig.). Ultimately, these changes resulted in control pattern and 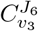 values that were indistinguishable from those of the reference model (S2 Fig.).

### Control patterns in the free-*ϕ*_*N*_ model

One final question that remains to be addressed concerns the use of the fixed-*ϕ*_*N*_ model (as reported up to now) instead of a model in which *ϕ*_*N*_ is a free variable and the activity of the existing NADH oxidase reaction (*υ*_13_) is modulated. S1 Table shows the dominant 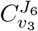 patterns in such a “free-*ϕ*_*N*_” model, which were chosen in the same manner as for the fixed *ϕ*_*N*_ model, expressed in terms of the 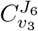 control patterns in the fixed-*ϕ*_*N*_ model. In all cases the free-*ϕ*_*N*_ control pattern expressions were very similar to those presented in Table 1, with most patterns differing only by two symbols when compared to their fixed-*ϕ*_*N*_ counterparts. Additionally, S4 Fig. shows the 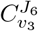 control patterns of the free-*ϕ*_*N*_ model in a similar manner to those of the fixed-*ϕ*_*N*_ model in Fig. 3. This demonstrates that while the value of 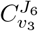 differed between the two models, the contributions of the control patterns towards the control coefficient responded very similarly towards changing *ϕ*_*N*_ values in spite of the differences between their control pattern expressions.

More convincing, perhaps, is that the flux responses of the free-*ϕ*_*N*_ model towards changing *ϕ*_*N*_ (as facilitated by the modulation of NADH oxidase activity) are exactly the same as those of the fixed-*ϕ*_*N*_ model (S5 Fig.). This indicates that while the control patterns are slightly altered by fixing the concentration of *ϕ*_*N*_ and varying it directly, this does not affect the overall flux and control behaviour of the system.

## Discussion

The results shown in this paper further explore those from a previous study [29] on a model of pyruvate-branch metabolism in *Lactococcus lactis* [21]. Surprisingly, we found that the flux through acetaldehyde dehydrogenase (*J*_6_) responded negatively towards an increase in the ratio of NADH to NAD^+^ (*ϕ*_*N*_), in spite of NADH being a substrate for this reaction; this was found to be the result of the interaction of NADH/NAD^+^ with pyruvate dehydrogenase (reaction 3) combined with its strong control over the *J*_6_ as quantified by 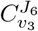 [29]. Thus, to ascertain what determines the value of 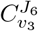 and how it is influenced by the properties of, and interactions between, the individual components of the pathway, we investigated this control coefficient using the frameworks of symbolic control analysis [12], control pattern analysis [14,38], and thermodynamic/kinetic analysis [16].

The algebraic expression for 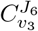 consisted of 76 control patterns, representing the totality of the chains of local effects that can potentially affect its ultimate value. However, only 11 of these patterns actually contributed significantly towards the total numeric value of 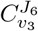 for the four orders of magnitude range of *ϕ*_*N*_ values investigated. Moreover, at most only six control patterns met our cut-off criteria for important contributors towards the control coefficient for any particular *ϕ*_*N*_ value. Over the full range of *ϕ*_*N*_, four different groups of control patterns were found to be dominant within different ranges of *ϕ*_*N*_ values with a clear change of “regime” from one group to the next as the *ϕ*_*N*_ value was varied. While the criteria for defining a control pattern as “important” were selected in order to minimise the number of control patterns and were very effective in our case, this strategy may not generally hold up in all systems. It is conceivable that the value of a control coefficient may be determined by a large group of control patterns where each contributes a small amount towards the total, instead of a small group contributing a large amount. Likewise, the remaining 85% of control patterns for 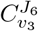 in our system could play a significant role under a different set of conditions; however, these conditions are unknown and were not investigated further.

Another strategy that decreased the number of control patterns was to fix the concentrations of NADH and NAD^+^, thus turning them into parameters of the system. While fixing NADH/NAD^+^ did have a minimal effect on control pattern composition, it nevertheless allowed us to simplify the analysis because the flux behaviour of the system was completely unaffected. The reason for this is that fixing the NADH/NAD^+^ ratio only removes the reaction catalysed by NADH oxidase from the system, and this reaction only has a single route of communication with the rest of the system (*viz.* NADH/NAD^+^). The effect that this reaction has in the free-NADH/NAD^+^ system is emulated in the fixed-NADH/NAD^+^ model by directly modulating NADH/NAD+. This, in turn, blocks communication between different parts of the pathway via NADH/NAD^+^. These two alterations in the structure of the system affect its control properties such that the summation property is not violated, i.e., the control coefficient values change but their sum remains equal to one. Communication between different parts of the system via routes other than those passing through NADH/NAD^+^ is unaffected. Note, however, that while this specific modification had relatively little effect on overall system behaviour, such an alteration will most probably yield significantly divergent results when choosing other intermediates that are not directly linked to a demand reaction such as NADH/NAD^+^ in this case. Thus, the strategy of fixing an internal metabolite for simplifying symbolic control analysis expression needs to be carefully considered.

Dividing the control patterns into their constituent backbone and multiplier patterns revealed that the control patterns within each of the four dominant groups were not only related in terms of behaviour and response towards *ϕ*_*N*_, but also in terms of composition. This makes intuitive sense as one would expect control patterns (and subpatterns) with similar composition to behave in a similar way. However, the influence of different control patterns on 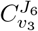 was determined by more nuanced factors than only their composition. In our system backbone patterns and T multiplier patterns determined the region in which control patterns were dominant, while B multipliers determined the magnitude of the control patterns within their group. Since these subpatterns correspond to actual metabolic branches within the pathway, it seems that each of these particular metabolic branches played a specific role in determining 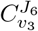. The general similarity of the different branches did not depend solely on their composition, e.g. when comparing B multiplier patterns, a similar response towards *ϕ*_*N*_ for each pattern was observed in spite of some of them not sharing any common flux or elasticity components. Thus, dissimilar control patterns can exhibit similar behaviour if they occur within the same metabolic branch. Divisions of control pattern factors along different branches could thus be a useful way for relating the control properties of a system to its network topology. Ultimately, while the concept of backbone and multiplier patterns was originally devised as a method for generating control coefficient expressions by hand [38], here it was extremely useful for simplifying the control pattern analysis by providing an easily digestible and descriptive language for comparing and contrasting control patterns. In addition, this methodology was indispensable for narrowing down the search for the lowest level metabolic components responsible for a particular high-level system behaviour.

Using the backbone and multiplier patterns as a starting point, we investigated two control patterns that were closely related in terms of composition (CP063 and CP071), but behaved differently because the expressions of their T multipliers differed in terms of two factors. These two control patterns were also representative of their control pattern groups (i.e. group 3 and group 4), since all patterns in the same group only differed by their B multipliers. By examining these factors as a function of *ϕ*_*N*_ it seemed that only the difference of a single factor actually contributed to the difference in observed behaviour on the control pattern level—i.e. the difference in sensitivity of the rates of the acetoin efflux step (reaction 10) and acetoin dehydrogenase reaction (reaction 11) towards acetoin concentration, as quantified by the elasticity coefficients 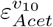 and 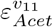 respectively. Since reaction 11 was modelled as a reversible step, 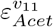 became massive as this reaction neared equilibrium, whereas 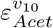 had a value of one, ultimately causing CP073 and all group 4 control patterns to have larger values at the highest tested *ϕ*_*N*_ values than CP063 and the group 3 patterns. These results suggest that seemingly insignificant components in a metabolic pathway can have remarkable effects on the ultimate control of the system. They also demonstrate that irreversible reactions can either have no effect on the magnitude of a control coefficient, or they can decrease its magnitude because the elasticity is limited to a range of values between zero and one. Conversely, reversible reactions can have an increasing, neutral, or decreasing effect on a control coefficient when accounting for both substrate and product elasticities and these elasticities can assume very large values as a reaction approaches equilibrium.

As discussed in our previous work [29], the observed decrease in flux towards ethanol (*J*_6_) in response to a large NADH/NAD^+^ value corresponds with previous observations that the NADH/NAD^+^ ratio plays a role in regulating the shift from mixed-acid fermentation to homolactic fermentation (e.g. [24,28]). While we previously established that 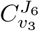 was a key component in causing the shift in the model, the current work expands on this by identifying the specific components responsible for determining 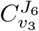, and thus the shift towards homolactic fermentation. Group 4 patterns (consisting of multiplier T6 and Backbone C) were responsible for the relatively large 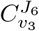 values within the same NADH/NAD^+^ range in which a decrease of *J*_6_ was previously observed. The large positive value of these control patterns can be attributed to three factors: first, the large magnitude of T6 relative to the other T multipliers, second, the large absolute values of the B multipliers, and third, the large absolute value of the C scaled backbone for NADH/NAD^+^> 0.0265, all of which act to increase the magnitudes of the control patterns in group 4 for the NADH/NAD^+^ range in which they are dominant. Thus, besides the fact that 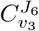 is just one of the factors responsible for the decrease in ethanol flux, the large value of 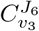 during the shift to homolactic fermentation can itself be seen as resulting from the interactions between numerous seemingly unrelated metabolic components. As will be reiterated below, this illustrates that metabolic engineering efforts must take into account much more than a few isolated components if large scale success is to be achieved.

Our results demonstrate that a great advantage of symbolic control analysis over numerical analysis of control coefficients is in its ability to provide a mechanistic explanation of control in terms of the low-level components of a system. This tool can thus truly be regarded as a systems biology framework [47]. In other work we have used symbolic control coefficient expressions as an explanatory tool [48] to analyse the control patterns responsible for the shift in control under different pH and environmental conditions in models of fermentation of *Saccharomyces cerevisiae* [49]. In that case an observed increase in glucose uptake rate in immobilised cells was shown to be the result of the activation of a subset of the system related to carbohydrate production. Symbolic control analysis was also used to mechanistically explain the ultrasensitive flux-response observed in a model of the *Escherichia coli* thioredoxin system [50]. In this work symbolic control coefficients expressions were not only useful for explaining ultrasensitivity, but also for deriving the quantitative conditions that need to be fulfilled for ultrasensitivity to occur.

Another advantage of symbolic control coefficient expressions over their numeric counterparts is that they rely only on knowledge of network topology and regulatory interactions. In other words, the same control coefficient (and control pattern) expressions hold true regardless of any particular steady-state conditions. Therefore the control of a system for which a full kinetic characterisation is unavailable can be predicted by substituting measured (or hypothetical) steady-state elasticity coefficient, flux, and concentration values into the symbolic control coefficient expressions to yield numeric control coefficient values. Similarly, this property also allows one to predict the control of any system under different conditions by substituting in elasticity coefficient values representing such conditions (e.g., reactions close to equilibrium, or irreversible reactions far from saturating concentrations). In contrast to our work, most of the past applications of symbolic control coefficient expressions have been of this type, e.g. [51, 52]. While symbolic control coefficient expressions are not strictly necessary to generate control coefficients from elasticity coefficients since numerical inversion of the **E**-matrix would achieve the same result, the relationship between these expressions and the structure of the metabolic pathway conveys more biological meaning and provides more granularity than a numerical matrix inversion. A recent example of such a treatment can be found in [53], where control of unregulated and feedback-regulated systems is compared using control coefficient expressions populated with hypothetical values. By demonstrating how different structures and conditions give rise to different system properties, the author not only predicts control from elasticity but explains these phenomena in terms of the system’s structure.

The thermodynamic/kinetic analysis framework complements symbolic control analysis and provides an additional layer of description by allowing the elasticity coefficients to be dissected into their binding and mass-action components. Since regulation of a reaction can be seen as the alteration of the effect of mass action (either through augmentation or counteraction) [17,54], this framework allows us to separate enzyme regulation from the properties of the the reaction itself. For instance, the counteraction of the effect of mass action on the sensitivity of reaction 11 towards acetoin by the effect of enzyme binding in the red shaded area of Fig. 4B indicates that the reaction would have been quite sensitive towards its substrate under the prevailing conditions had it not been for the presence of the enzyme (although it would have occurred at a lower rate). Additionally, while not explored in detail in this paper, this framework also allows for the separation of the effects of kinetics and thermodynamics, and allows for the quantification of the effects allosterism and cooperativity [16]. The use of other approaches that split rate equations into smaller components (thus highlighting different effects) [55,56], may also prove to be useful when combined with symbolic control analysis.

While we were able to describe system behaviour in terms of component behaviour, an attempt to use this information to modify the control properties of the system via an alteration of the properties of reaction 11 proved to be fruitless. By effectively making the reaction irreversible, we hoped to decrease the magnitude of 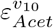 to cause CP071 to have a smaller value than CP063. While this change did alter 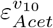, the system almost completely compensated for the change, resulting in no practical difference between the control properties of the reference and altered systems. This illustrates that while symbolic control analysis is an excellent tool for understanding control in mechanistic terms, it does not necessarily yield easy answers for use in metabolic engineering due to the overall complexity of metabolic systems and their ability to adapt to changes. It does however allow us to investigate such hypothetical manipulations of the system with relative ease and, in our case, it demonstrated the homeostatic properties of the system. Thus, while it is tempting to fully ascribe system properties to single metabolic components, one must be careful not to fall into the trap of viewing metabolic systems in a reductionistic manner. Nevertheless, symbolic control analysis may indeed have the potential to be used in metabolic engineering, but it will require a more nuanced approach than the one demonstrated here.

The analysis presented in this paper expanded on previous work by delving deeper into one of the causes for the observed shift away from ethanol production at high NADH/NAD^+^ values using the frameworks of symbolic control analysis and thermodynamic/kinetic analysis. The detailed mechanistic description and analysis of the control coefficient responsible for the large shift provided new insight into this phenomenon. Additionally, this work represents the first of its kind to analyse control of a realistic metabolic model on such a low level by using the concepts of control patterns and their constituent backbone and multiplier patterns. Our hypothetical manipulation of the system based on our new-found knowledge also reiterated the danger of viewing metabolic systems from an overly reductionistic perspective and highlight the need for more robust metabolic engineering strategies. While we believe that the techniques used in this paper are mostly suited to describing and understanding the behaviour of metabolic systems in a particular steady state, such understanding is an important stepping stone to developing practical methods for manipulating metabolic control.

## Acknowledgements

This work was supported by the National Research Foundation of South Africa (NRF) [81129 to J.M.R., 84611 to C.D.C.].

## Supporting information

**S1 Example analysis** Jupyter notebook, additional code required, model description files and instructions to recreate the computational analysis.

**S1 Fig.** The flux and elasticity components of T4 and T6 as functions of *ϕ*_*N*_ after alteration of *K*_*eq*_ of reaction 11.

**S2 Fig.** The most important control patterns of 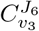 as functions *ϕ*_*N*_ of after alteration of *K*_*eq*_ of reaction 11.

**S3 Fig.** Backbone and multiplier patterns of the 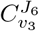 control patterns as functions of *ϕ*_*N*_ after alteration of *K*_*eq*_ of reaction 11.

**S4 Fig.** The most important control patterns of 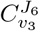 as functions of *ϕ*_*N*_ in the free-NADH/NAD^+^ model.

**S5 Fig.** Rate characteristic plots of the reaction block fluxes of *ϕ*_*N*_.

**S1 Table** Numerator expressions of the dominant control patterns of 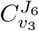 in the free-NADH/NAD^+^ model.

